# Accumulation of mitochondrial ROS drives reductive glutamine metabolism

**DOI:** 10.1101/2022.09.15.508079

**Authors:** Xiangpeng Sheng

## Abstract

Hypoxia, mitochondrial defect or extracellular matrix detachment induces reductive glutamine metabolism that can produce citrate by carboxylating the glutamine-derived ketoglutarate (α-KG). Reductive carboxylation is required for maintaining redox homeostasis, lipogenesis and cancer cell growth. However, the fundamental mechanism(s) of how reductive glutamine metabolism may be controlled is not fully understood. This study here demonstrates that mitochondrial-derived reactive oxygen species (ROS) positively drives reductive carboxylation. Knockout of fatty acid synthase (FASN) causes increased mitochondrial ROS and significant reductive glutamine metabolism. Mitochondrial ROS accumulation is identified as a shared feature in cells that utilize reductive metabolism. Moreover, ROS scavenger could inhibit reductive carboxylation by decreasing ROS level in mitochondria. By contrast, ROS inducers activate reductive carboxylation via increasing mitochondrial ROS. Mitochondrial ROS level determines the degree of reductive carboxylation with a positive correlation. The results presented here not only reveal a fundamental mechanism of how reductive carboxylation is controlled, but also elucidate a critical role of ROS in reprogramming metabolism.

## 1. Introduction

Mitochondrial oxidative metabolism generates intermediates that support the macromolecular biosynthesis as well as the cell proliferation, whereas environmental or genetic interventions may trigger metabolic reprogramming [1, 2]. Glutamine, a major nutrient required for cancer cell growth, can be metabolized by the oxidative TCA cycle or reductive metabolism pathway [3, 4]. Reductive glutamine metabolism is characterized by a reductive carboxylation reaction controlled by NADPH-dependent isocitrate dehydrogenases (IDH) [5–7]. NADPH/CO2-dependent reductive carboxylation of αKG to isocitrate can be catalyzed by IDH1 and IDH2, two of three isoforms of IDH [5, 7]. Recently, reductive metabolism has been identified as a major carbon source for lipid synthesis and tumorigenesis during hypoxia, mitochondrial defect or extracellular matrix detachment [5-81. Since alterations in the carbon source selection may sustain the uncontrolled cell growth [6, 8], understanding this metabolic switch is of particular importance towards the development of promising cancer therapies [3, 9].

Because α-KG is a direct participant of reductive carboxylation reaction, α-KG alterations undoubtedly affect this process. Recent reports have suggested that α-KG oxidation and α-KG /citrate ratio have been reported to regulate reductive glutamine metabolism [9, 10]. Moreover, several particular proteins might be involved in the utilization of reductively metabolized glutamine [11–13]. However, the fundamental mechanism(s) of how hypoxia, mitochondrial impairment or matrix detachment activates glutamine-dependent reductive carboxylation is not fully understood. Therefore, by using metabolomics and stable isotope tracing, this study here aims to identify the common mechanism(s) of how reductive carboxylation is biochemically operated in cells.

## 2. Results

### 2.1. Knockout of fatty acid synthase (FASN) activates reductive carboxylation

Given the prioritized use of reductively metabolized glutamine during lipogenesis [5], blockade of lipid synthesis by inhibiting FASN may potentially trigger metabolic changes. Hence, FASN knockout clones of lung cancer H460 cells were presently constructed using the CRISPR/Cas9 technology (Fig. 1A and Fig. S1). To investigate the effects of FASN inhibition in mitochondrial metabolism, both wild-type (WT) and FASN knockout (KO) cell lines were cultured in medium containing uniformly labelled [U-^13^C]glucose, and ^13^C-enriched intracellular metabolites were then quantified by mass spectrometry (Fig. 1B). In WT H460 cells, most citrate molecules contained one or more glucose-derived ^13^C (Fig. 1C and Fig. S2A). However, unlabeled citrate was significantly increased in FASN KO cells, indicating that FASN deficiency may suppress the entry of glucose-derived carbon into the citrate structure (Fig. 1C).

**Fig. 1.**
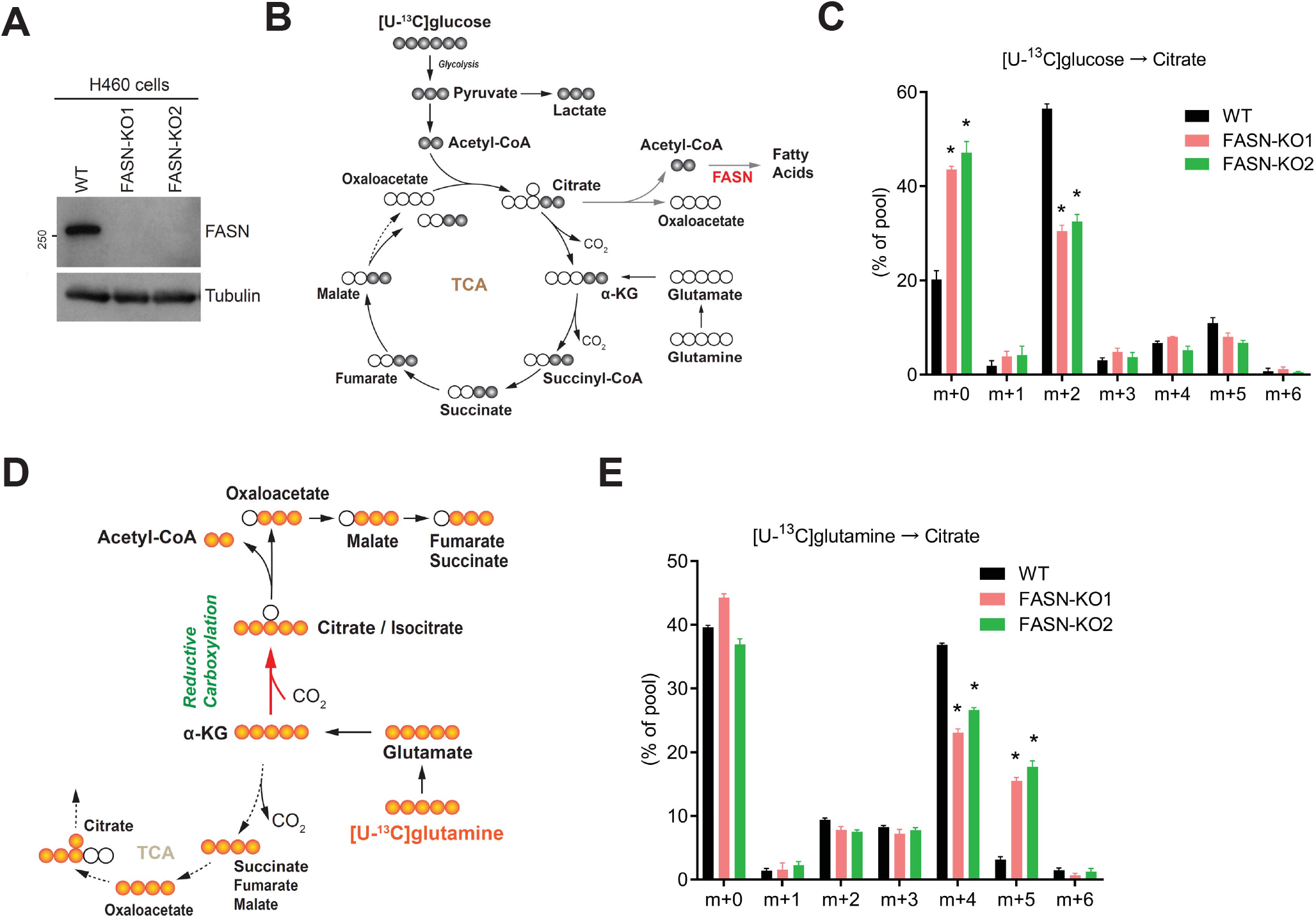
FASN deficiency impairs mitochondrial oxidative metabolism but activates reductive carboxylation. **(A)** Western blot analysis of FASN protein in WT or FASN KO H460 cell lines. **(B)** Schematic depicting the mitochondrial oxidative metabolism for citrate production from [U-^13^C]glucose. **(C)** Mass isotopomer analysis of citrate in WT or FASN KO cell lines cultured with [U-^13^C]glucose and unlabelled glutamine for 4 hours. \**P* < 0.05 (*n* = 3 independent cultures). **(D)** Schematic showing the reductive and oxidative glutamine metabolism for citrate production from [U-^13^C]glutamine. **(E)** Mass isotopomer distribution of citrate in WT or FASN KO cell lines cultured with [U-^13^C]glutamine and unlabelled glucose for 4 hours. \**P* < 0.05 (*n* = 3 independent cultures). All data represent mean ± SD, and *P* values were established using Student’s *t*-test.

To estimate the contribution of oxidative and reductive glutamine metabolism in the citrate synthesis, WT and two FASN KO cell lines were further cultured with [U-^13^C]glutamine. Labelled glutamine could either enter the oxidative pathway to produce citrate m+4 (i.e. citrate containing four ^13^C) or be directed towards a reductive pathway to generate citrate m+5 (Fig. 1D). All cell lines showed some comparable ^13^C labelling of the total citrate pool, but the overall ^13^C distribution was different between WT and KO cells (Fig. 1E and Fig. S2B). WT H460 cells used glutamine to produce a large amount of citrate m+4 and only limited quantities of citrate m+5 (~3%). In contrast, the citrate m+5 was significantly increased (15%) while citrate m+4 was decreased in FASN KO cells (Fig. 1E). These data suggest that FASN deletion may impair the mitochondrial oxidative metabolism but, at the same time, it can promote reductive carboxylation.

### 2.2. Mitochondrial ROS accumulation in FASN-KO cells is required for reductive metabolism

These intriguing findings prompted me to explore how FASN deletion may activate reductive carboxylation. Pathological effectors such as hypoxia, mitochondrial impairment and matrix detachment are three well-established inducers of reductive carboxylation in cells [5–8]. Some common cellular features based on reductive carboxylation and reactive oxygen species (ROS) accumulation in mitochondrial have been noticed. In fact, hypoxia is known to trigger robust mitochondrial ROS generation [14, 15], while dysfunctional mitochondria can serve as a signaling platform to promote ROS accumulation [16, 17]. Moreover, matrix detachment has been shown to induce a significant increase in ROS levels [18]. Therefore, I hypothesize that reductive carboxylation is accompanied by mitochondrial ROS accumulation.

Since active reductive carboxylation is induced by FASN depletion, an increase in ROS production in FASN KO cells may potentially occur. Indeed, FASN KO cells presented a higher mitochondrial ROS level relative to WT H460 cells (Fig. 2A and B). Moreover, treatment with N-acetyl-L-cysteine (NAC), a direct ROS scavenger, was able to reduce mitochondrial ROS efficiently in both WT and KO cells (Fig. 2A and B). Intriguingly, a significant decrease in reductive carboxylation activity was detected in all NAC-related groups (Fig. 2C), suggesting that NAC-mediated ROS reduction impairs reductive carboxylation. Since NADPH directly participates in reductive metabolism [8], the NADPH/NADP^+^ ratio was further examined. Although the NADPH/NADP^+^ ratio was reduced in FASN KO cells, NAC treatment had no effect on this proportion (Fig. 2D), indicating NADPH is not involved in NAC-mediated inhibition of reductive carboxylation. Thus, increased mitochondrial ROS levels appear to be responsible for activating reductive carboxylation in FASN KO cells.

**Fig. 2.**
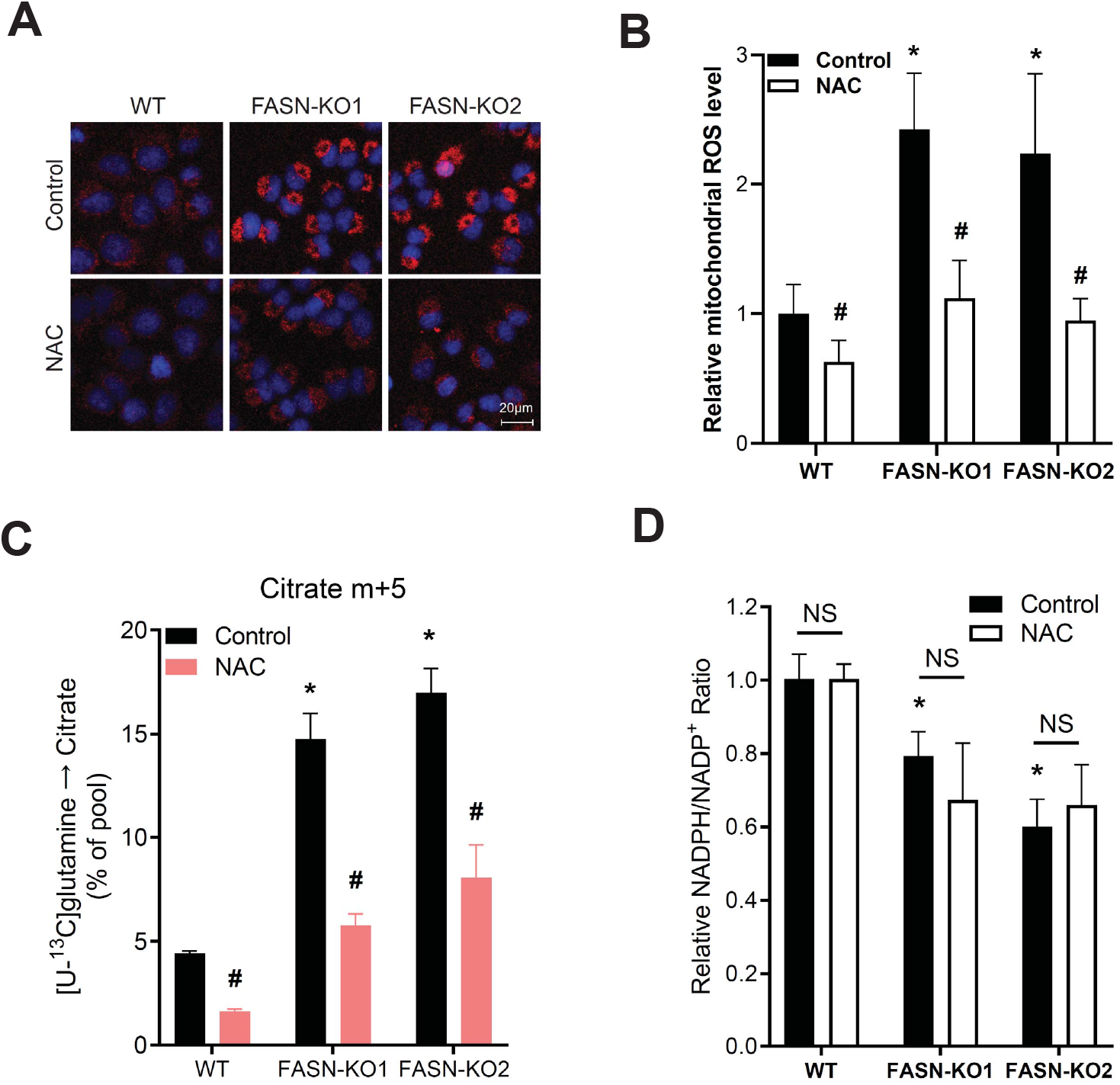
Accumulated mitochondrial ROS induces reductive metabolism in FASN KO cells. **(A**) Mitochondrial ROS was measured using MitoSOX staining. WT and FASN KO cell lines were treated with or without ROS scavenger NAC (10 mM, 4 h). Representative images of MitoSOX staining (red), Hoechst staining of cell nuclei (blue) are shown (Scale bar, 20 μm). **(B)** Statistical results of relative ROS level in **(A)**. \**P* < 0.05 (KO1 or KO2 versus WT), #*P* < 0.05 (NAC versus Control) (*n* = 50 cells). **(C)** Abundance of citrate m+5 in cells cultured with [U-^13^C]glutamine and treated with or without NAC. The labelled glutamine was introduced at the same time the cells were treated with or without NAC. Metabolites were extracted 4 hours later. \**P* < 0.05 (KO1 or KO2 versus WT), #*P* < 0.05 (NAC versus Control) (*n* = 3 independent cultures). **(D)** NADPH/NADP^+^ ratio was measured in WT or FASN KO H460 cells treated with or without NAC. \**P* < 0.05 (KO1 or KO2 versus WT), NS represents not significant (*n* = 3 biological repeats). All data represent mean ± SD, and *P* values were established using Student’s *t*-test.

### 2.3. Mitochondrial ROS inducer can activate reductive glutamine metabolism

Further investigations were pursued to verify whether other mitochondrial ROS inducer(s) could also activate reductive carboxylation. Dehydroepiandrosterone (DHEA), a pentose phosphate pathway (PPP) inhibitor, has been shown to markedly enhance mitochondrial ROS [7, 19]. Therefore, H460 cells were properly treated with DHEA and, as expected, this reagent was capable of inducing a significant increase in mitochondrial ROS as well as reductive carboxylation activity in cells (Fig. 3A and B). On the other hand, treatment with the anti-oxidant NAC not only decreased mitochondrial ROS accumulation but also inhibited DHEA-induced reductive carboxylation (Fig. 3C). Furthermore, DHEA treatment was able to reduce the NADPH/NADP^+^ ratio, but this proportion was not affected by NAC (Fig. 3D), thus indicating that NAC regulates reductive carboxylation by decreasing ROS levels but not altering NADPH content. These data suggest that the accumulation of mitochondrial ROS drives reductive carboxylation in cancer cells.

**Fig. 3.**
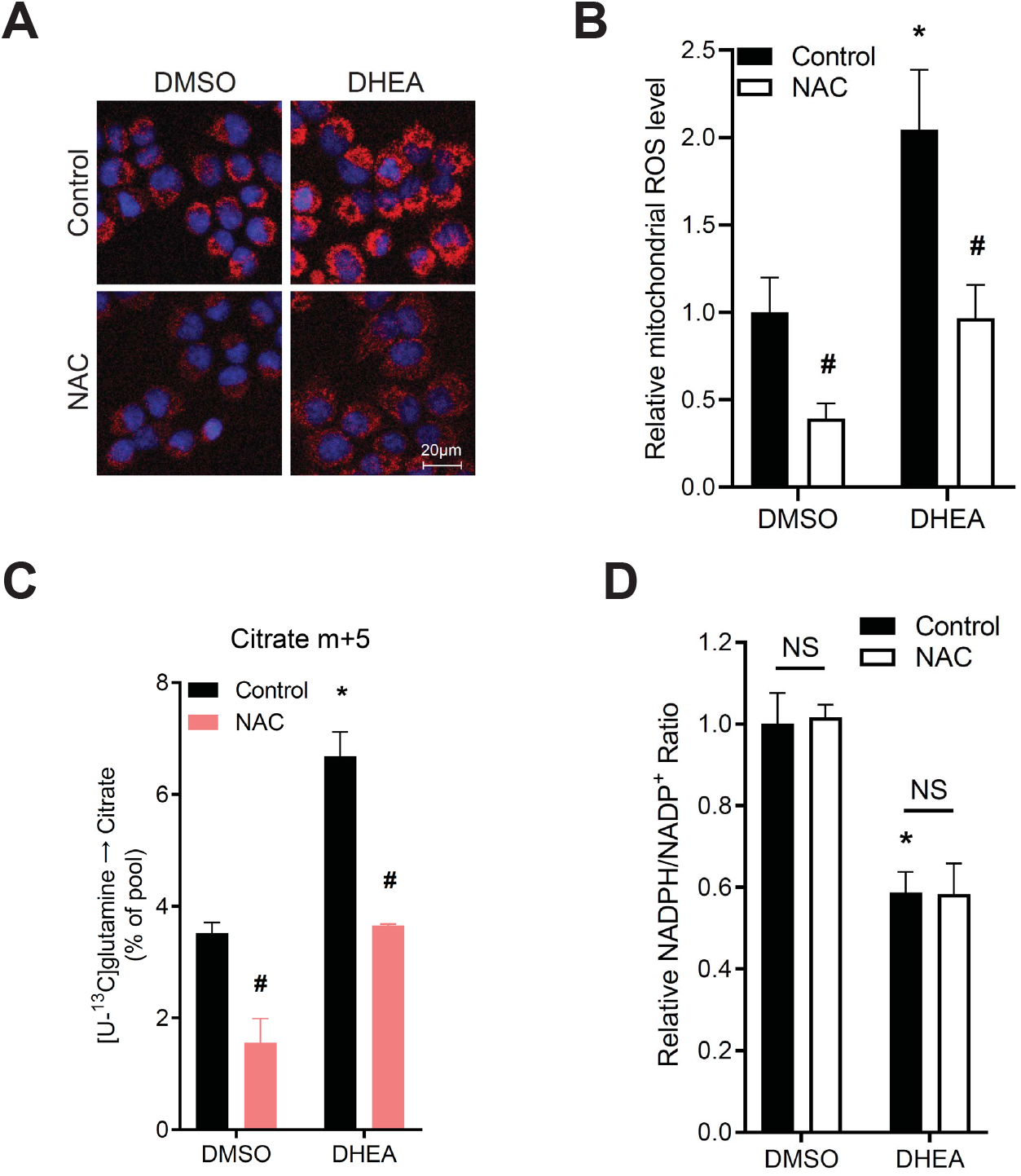
Mitochondrial ROS inducer DHEA activates reductive carboxylation. **(A**) Mitochondrial ROS was measured in DHEA-treated WT H460 cells using MitoSOX staining (red) and Hoechst staining (blue). Cells were treated with DHEA (100 uM, 24 h) and/or NAC (10 mM, 4 h). The representative images are shown. Scale bar, 20 μm. **(B)** Statistical analysis of mitochondrial ROS levels in (A). \**P* < 0.05 (DHEA versus DMSO), #*P* < 0.05 (NAC versus Control) (*n* = 50 cells). **(C)** Abundance of citrate m+5 from [U-^13^C]glutamine in WT H460 cells with indicated treatment. Cells were pre-treated with DHEA (100 μM, 20 h) in a regular medium, and were further treated with NAC in [U-^13^C]glutamine medium for 4 hours. \**P* < 0.05 (DHEA versus DMSO), #*P* < 0.05 (NAC versus Control) (*n* = 3 independent cultures). **(D)** NADPH/NADP^+^ ratio was measured in WT H460 cells with DHEA and/or NAC treatment. \**P* < 0.05 (DHEA versus DMSO), NS represents not significant (*n* = 3 biological repeats). All data represent mean ± SD, and *P* values were established using Student’s *t*-test.

Exogenous H_2_O_2_ treatment can induce intracellular oxidative stress and cell death [20, 21], suggesting exogenous H_2_O_2_ treatment may promote reductive glutamine metabolism. In H460 cells, exogenous H_2_O_2_ readily induced cell death (Fig. S3A). However, no significant alterations were observed in the proportions of citrate m+5 from [U-^13^C]glutamine after H_2_O_2_ treatment (Fig. S3B). Of note, although mitochondrial ROS level was increased in the early period (1 hour) of H_2_O_2_ treatment, the increasement of mitochondrial ROS disappeared completely after 2 or 4 hours (Fig. S3C), consistent with the invariable level of reductive carboxylation. These data further demonstrate a requirement of ROS accumulation in mitochondria for promoting reductive carboxylation.

### 2.4. Mitochondrial ROS level positively determines the degree of reductive carboxylation

Since data related to NAC treatment have shown that reducing ROS levels could lower reductive carboxylation activity, I hypothesize that a higher mitochondria ROS content can potentially drive some stronger reductive metabolism. To test this hypothesis, H460 cells were treated with Diphenyliodonium (DPI), an alternate ROS inducer in some cells [19, 22]. It was observed that DPI could induce much higher ROS levels in mitochondria when compared to DHEA (Fig. 4A). Consistently, reductive carboxylation was also stronger in DP1 than DHEA groups (Fig. 4B). Moreover, upon treatment of FASN KO cells with DHEA, higher mitochondrial ROS level as well as stronger reductive carboxylation were observed, showing an additive effect between FASN knockout and single DHEA treatment (Fig. 4C, D and E). These data demonstrate that mitochondrial ROS level is positively correlated with the degree of reductive carboxylation.

**Fig. 4.**
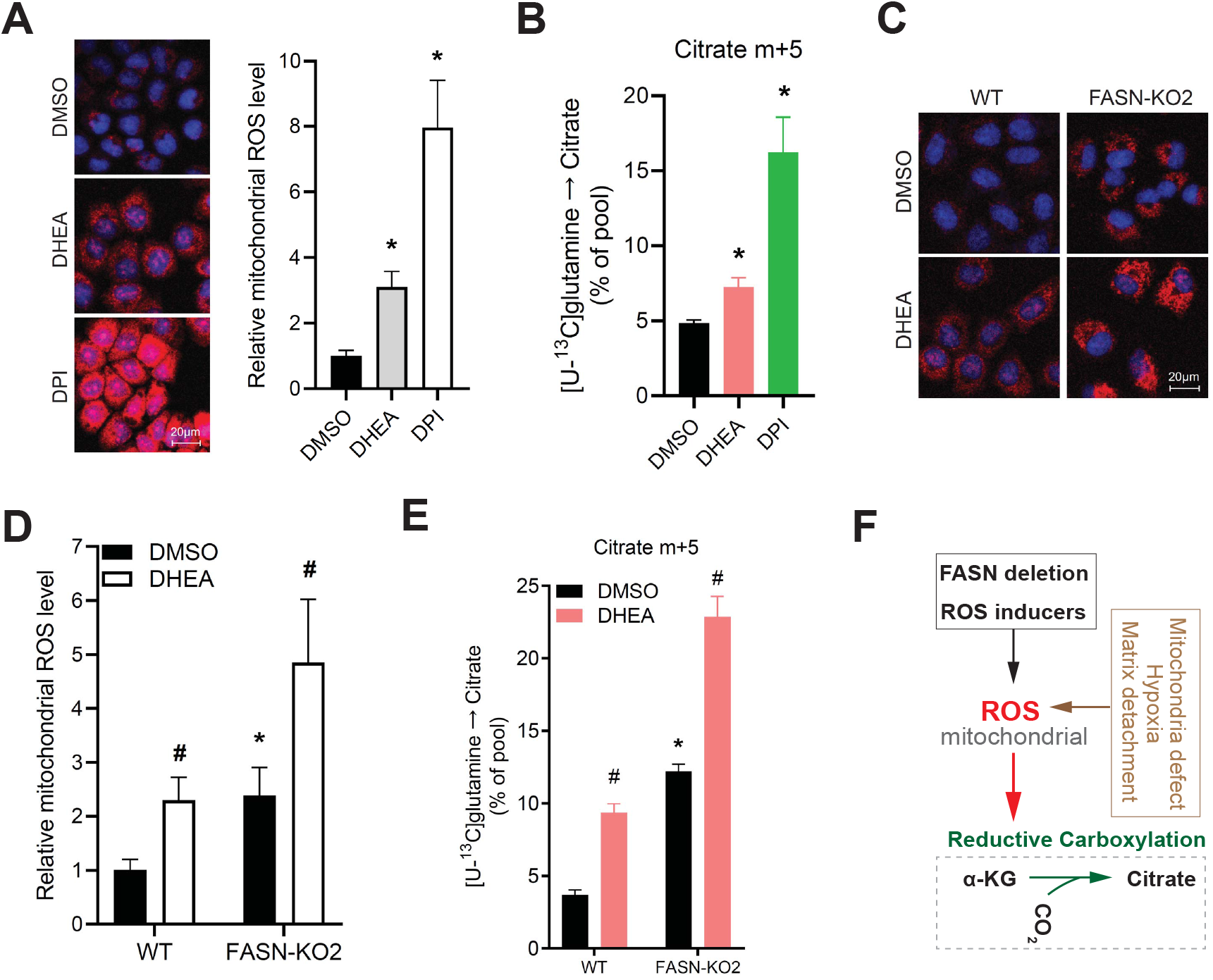
The degree of reductive metabolism is determined by mitochondrial ROS level. **(A)** Mitochondrial ROS was determined in H460 cells treated with ROS inducers. Treatment of WT H460 cells with DHEA (100 μM, 24 hours) or DPI (5 μM, 4 hours). MitoSOX staining (red) showed the mitochondrial ROS and Hoechst (blue) indicated the cell nuclei. Representative images and ROS quantitation are shown. Scale bar, 20 μm. \**P* < 0.05, (*n* = 50 cells). **(B)** Mass isotopomer analysis for citrate m+5 from WT cells cultured in medium containing [U-^13^C]glutamine and ROS inducers for 4 hours. \**P* < 0.05 (*n* = 3 independent cultures). **(C)** MitoSOX staining (ROS, red) and Hoechst staining (nuclei, blue) in DHEA-treated WT or FASN KO H460 cells. Representative images are shown. Scale bar, 20 μm. **(D)** ROS quantitation in **(C)**are shown. \**P* < 0.05 (KO2 versus WT), #*P* < 0.05 (DHEA versus DMSO) (*n* = 50 cells). **(E)** Abundance of citrate m+5 in WT or FASN KO H460 cells cultured with [U-^13^C]glutamine and treated with DMSO or DHEA. \**P* < 0.05 (KO2 versus WT), #*P* < 0.05 (DHEA versus DMSO) (*n* = 3 independent cultures). **(F)** A schematic showing that FASN deficiency, ROS inducers or other pathological factors can activate reductive carboxylation via inducing mitochondrial ROS accumulation. All data represent mean ± SD, and *P* values were established using Student’s *t*-test.

## 3. Discussion

Collectively, the current study clearly indicates that mitochondrial ROS positively drives reductive glutamine metabolism. FASN knockout (or ROS inducers) induced the accumulation of mitochondrial ROS and an active reductive carboxylation in cancer cells (Fig. 4F). Moreover, ROS scavenger exhibited a negative regulation towards reductive metabolism by decreasing mitochondrial ROS level. Thus, reductive carboxylation could be manipulated by controlling mitochondrial ROS in cells.

Mitochondria is a major source of ROS in cells [23]. Notably, cancer cells generate increased ROS relative to normal cells [24, 25]. Because increased mitochondrial ROS acts as signaling molecules to promote cancer cell proliferation and tumorigenesis, targeting mitochondrial ROS is an attractive therapy for some cancers [17, 26, 27]. The results here have demonstrated a novel and critical role of mitochondrial ROS to regulate reductive carboxylation in cancer cells, suggesting that the pro-growth responses of mitochondrial ROS may need active reductive glutamine metabolism.

As mentioned above, ROS accumulation in mitochondria is a common feature in cells that have active reductive metabolism, specifically during hypoxia, mitochondrial defect or matrix detachment. Of note, the retinal pigment epithelium also has mitochondrial oxidative stress and exhibits an high capacity for reductive carboxylation [21, 28], consistent to the concept of the current study here. The real connection between mitochondrial ROS level and reductive carboxylation in the retinal pigment epithelium needs to be elucidated by further investigations.

Lastly, future studies are warranted to elucidate the molecular mechanisms of how the increased mitochondrial ROS promotes reductive carboxylation. One possible mechanism is that the increased ROS in mitochondria might regulate the expression or enzymatic activities of some proteins associated to mitochondrial metabolism in cancer cells.

## 4. Materials and methods

### 4.1. Reagent

Anti-FASN rabbit antibody (#3189) was purchased from Cell Signaling Technology (CST). MitoSOX Red mitochondrial superoxide indicator (for live-cell imaging) (M36008) and Hoechst 33342 (H3570) were purchased from Invitrogen. Diphenyleneiodonium chloride (DPI, D2926) and N-acetyl-L-cysteine (NAC, A9165) was purchased from Sigma. Dehydroisoandrosterone (DHEA, AC154980025) was from ACROS Organics™. NADP/NADPH-Glo™ Assay kit (G9081) was from Promega.

### 4.2. Cell culture and transfection

H460 cells were grown in RPMI medium supplemented with 5% fetal bovine serum (FBS), 4 mM L-glutamine, 100 units/ml penicillin and 100 mg/ml streptomycin (HyClone). H460 Cells were cultured in a humidified atmosphere with 5% CO_2_ at 37 °C. Transfection was performed with Lipofectamine 2000 (Invitrogen) according to the manufacturer’s instruction.

### 4.3. Construction of FASN knockout cell line

FASN knockout H460 cell lines were constructed using CRISPR/Cas9 system[29]. Negative sgRNA and targeting sgRNA were separately inserted into pSpCas9(BB)-2A-GFP(PX458) (Addgene ID: 48138). Plasmid was transfected into H460 cells for 48 hours. After transfection, GFP-positive cells were sorted into 96-well plates at cell density of single cell/well. Cells were cultured for two weeks. Expression of FASN in each clone was determined by western blot analysis. To minimize variations among individual clones, three wild-type single clones were pooled together as WT cell line and eight knockout clones were pooled separately as two FASN KO cell lines.

### 4.4. Mass spectrometry-based metabolite profiling

Mass isotopomer analysis of metabolites using Gas Chromatography Mass Spectrometry (GC-MS) approach was performed as described before with slight modifications[7, 8]. Uniformly labelled [U-13C]glucose and [U-^13^C]glutamine were purchased from Cambridge Isotope Laboratories. All stable isotope tracing studies for H460 cells were performed in medium containing 5% dialyzed FBS and 1% HEPES. DMEM without glucose and glutamine was prepared from powder (Sigma-Aldrich, D5030) and supplemented with either 10 mM [U-^13^C]glucose and 3 mM unlabeled glutamine, or 10 mM unlabelled glucose and 4 mM [U-^13^C]glutamine.

Cells were cultured in 6 cm dishes until 80% confluent. Old medium was completely removed and then cells were incubated in fresh ^13^C-containing medium for 4 hours at 37 °C. Cells were washed with cold NaCl solution and were collected in 900uL of a cold 1:1 mixture of methanol and water, and then subjected to freeze-thawing cycle for three times. After centrifugation (12,000 rpm for 15 min at 4 °C), the supernatants with aqueous metabolites were collected to a new tube and evaporated completely under airflow at room temperature, while the pellets were dissolved in 500uL NaOH (100 mM) at 100 °C for protein concentration analysis. Next, the evaporated metabolites were dissolved in 50 μL methoxyamine hydrochloride (Sigma) at 42 °C for 1 hour and then derivatized with 100 μL tert-butyldimethylsilyl (TBDMS) for 90 min at 72 °C. An Agilent 7890B gas chromatograph connected to an Agilent 5977B mass selective detector were used to analyze metabolizes. Mass isotopomer distributions in each metabolite were determined by analyzing the appropriate ion fragments using MATLAB software (MathWorks), integrating the abundance of all mass isotopomers from m+0 to m+n, where m is the mass of the fragment ion without any ^13^C and n is the number of carbons in this metabolite. Finally, the abundance of each mass isotopomer was converted into a percentage of the total pool.

### 4.5. Mitochondrial ROS detection

Mitochondrial ROS was measured using MitoSOX™ Red mitochondrial superoxide indicator (for live-cell imaging) (M36008, Invitrogen). An appropriate volume of MitoSOX™ working solution (5 μM) and Hoechst 33342 (5 μg/mL) was added to cover cells adhering to coverslip(s). Cells were incubated at 37°C for 10 min and cells were protected from light. Cells were then washed gently with warm medium for three times and maintained in warm medium for confocal imaging (ZEISS LSM 700). Relative ROS levels were calculated using Photoshop (Adobe).

### 4.6. NADPH/NADP+ quantitation

NADPH and NADP^+^ were measured individually using NADP/NADPH-Glo™ Assay kit (G9081, Promega) according to the manufacturer’s instruction and the relative NADPH/NADP^+^ ratio was calculated.

### 4.7 Western blotting

Cells were washed with ice-cold PBS and lysed in RIPA buffer. Proteins were separated using SDS-PAGE and transferred to a nitrocellulose membrane. After blocking in 5% BSA, membranes were incubated with Anti-FASN rabbit antibody (#3189, CST) antibody or mouse anti-tubulin antibody (Sigma). Protein was next incubated with HRP-conjugated secondary antibodies and was detected using chemiluminescence.

#### Statistical analysis

Data were analyzed using either Microsoft Excel or GraphPad Prism 8. Statistical significance of the data was established using Student’s *t*-test, and *P* < 0.05 was considered statistically significant.

## Author contributions

X.S. designed the project, performed all the experiments and data analysis, and wrote the manuscript.

## Supplementary Figures

**Figure S1.**
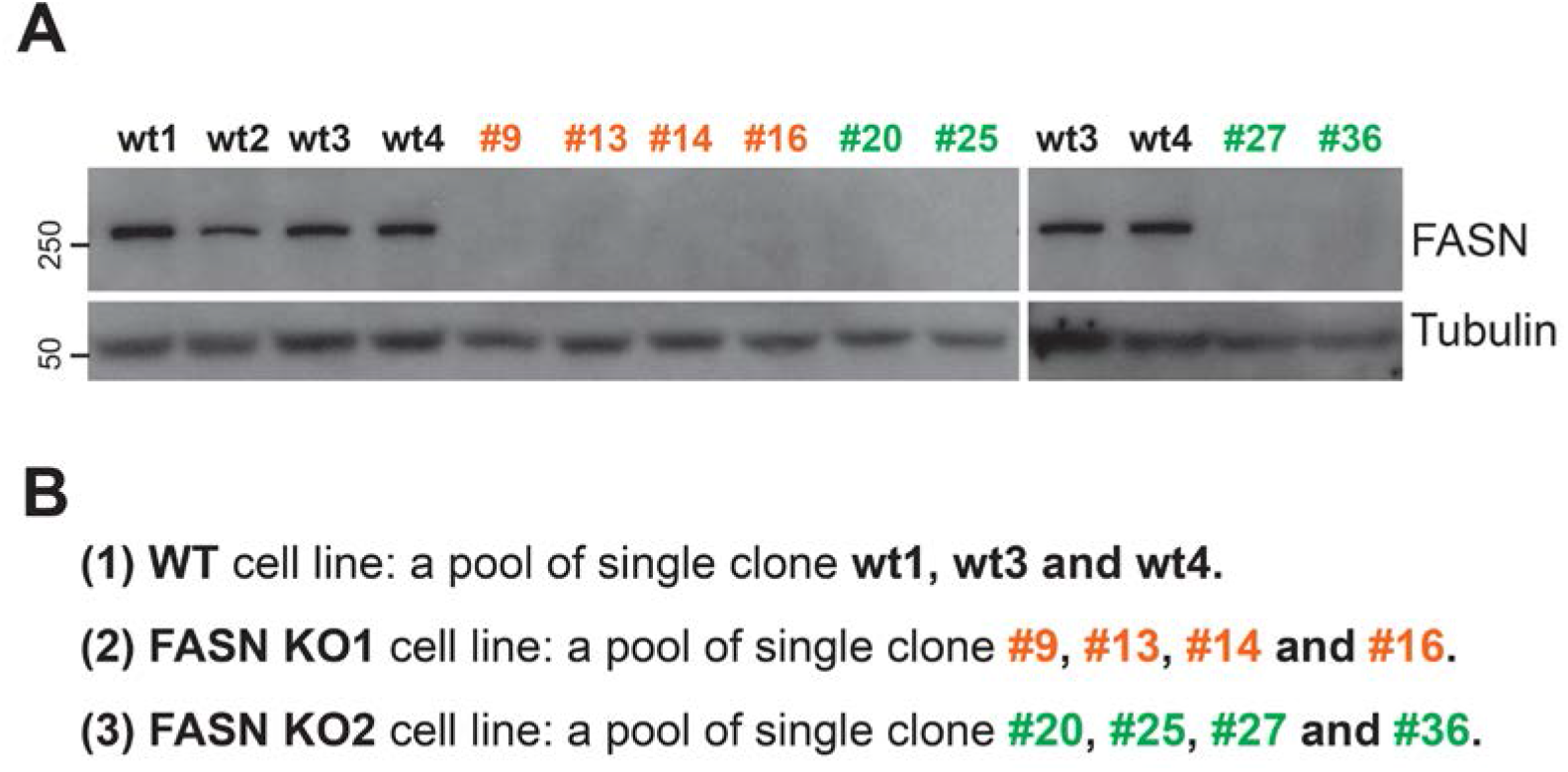
Construction of FASN knockout cell lines. **(A)** Western blot analysis of FASN expression in H460 cell clones. pSpCas9(BB)-2A-GFP(PX458) carrying negative or targeting sgRNA was transfected into H460 cells for 48 hours. GFP-positive cells were sorted into 96-well plates with single cell per well. After two weeks, cell clones were pickup for expansion and western blot analysis. **(B)** Construction of WT, FASN-KO1 and FASN KO2 cell lines by pooling several single clones together to control possible variations among individual clones.

**Figure S2.**
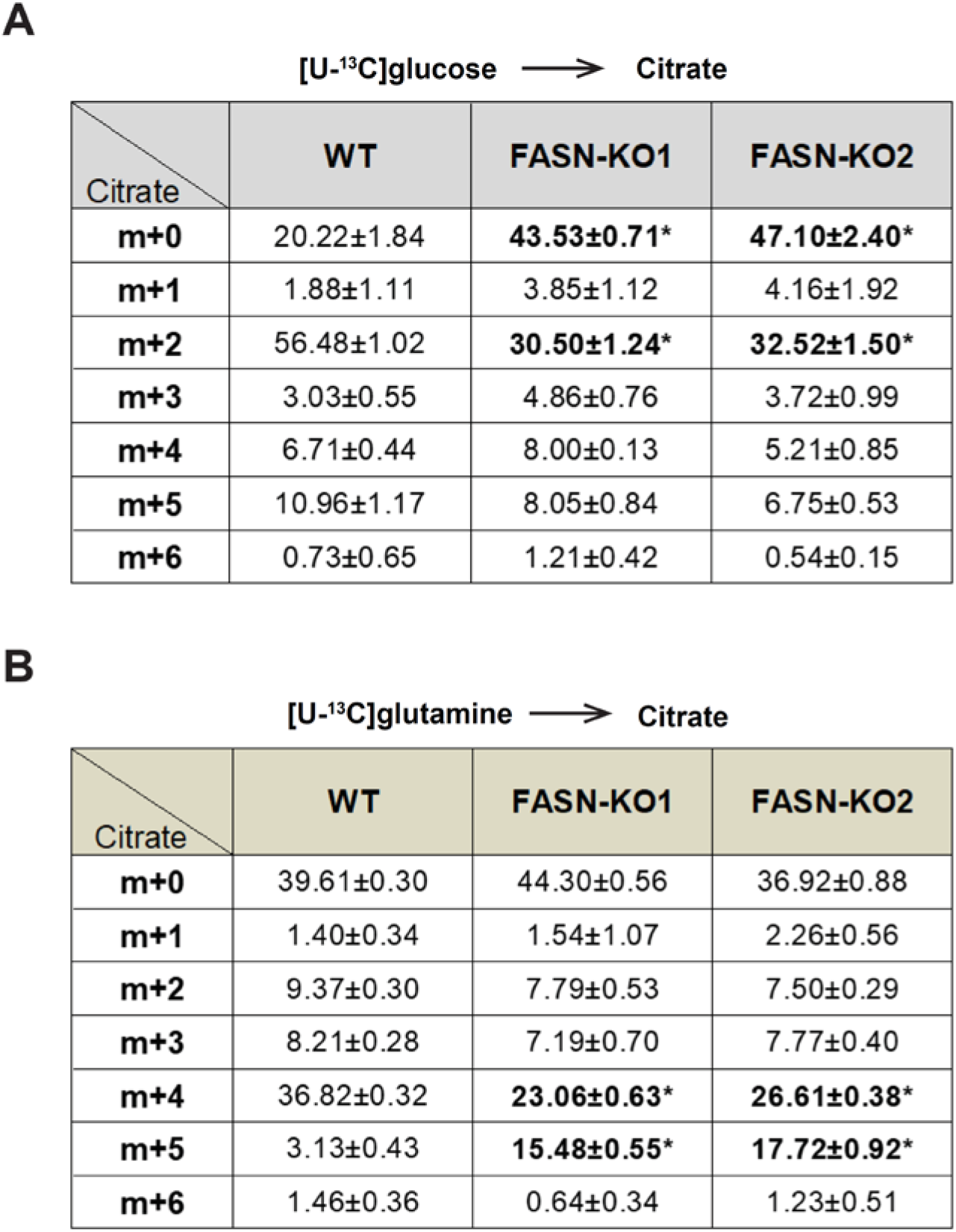
FASN knockout suppresses TCA cycle but promotes reductive carboxylation. **(A)** Mass isotopomer analysis of citrate in WT and FASN KO cell lines cultured with [U-^13^C]glucose and unlabelled glutamine for 4 hours. Data represent mean ± SD, \**P* < 0.05 (Student’s *t*-test, *n* = 3 independent cultures). Related to Fig. 1C. **(B)** Mass isotopomer distribution of citrate in WT and KO cell lines cultured with [U-^13^C]glutamine and unlabelled glucose for 4 hours. Data represent mean ± SD, \**P* < 0.05 (Student’s *t*-test, *n* = 3 independent cultures). Related to Fig. 1E.

**Figure S3.**
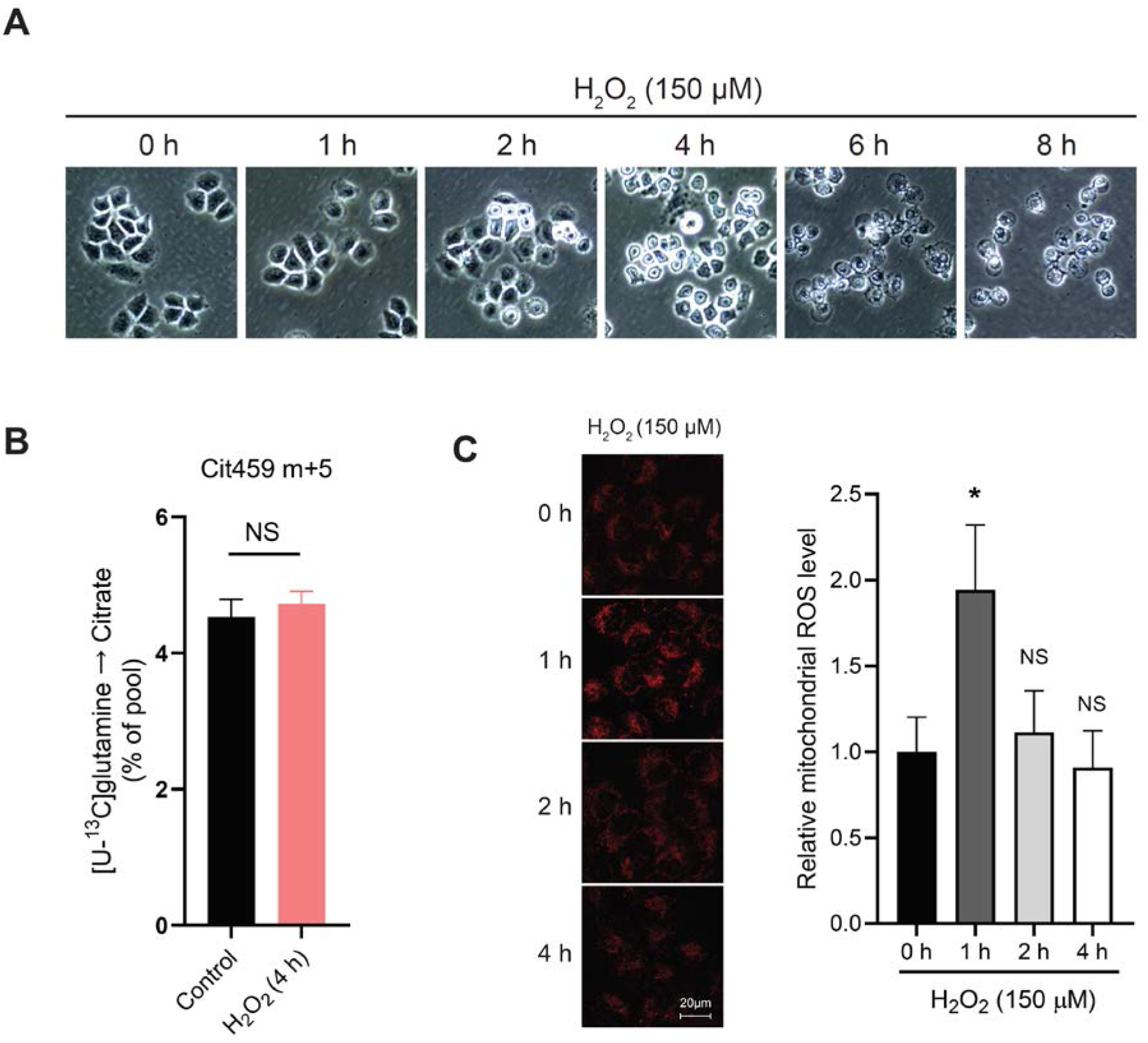
Exogenous H_2_O_2_ treatment in WT H460 cells. **(A)** WT H460 cells were treated with 150 μM H_2_O_2_. Cells were imaged at indicated time points. **(B)** Abundance of citrate m+5 from [U-^13^C]glutamine in H460 cells with or without H_2_O_2_ treatment. Cells were treated with 150 μM H_2_O_2_ in [U-^13^C]glutamine medium for 4 hours. NS represents not significant (*n* = 3 independent cultures). **(C)** Mitochondrial ROS was determined in H460 cells treated with H_2_O_2_. MitoSOX staining (red) showed the mitochondrial ROS. Representative images and ROS quantitation are shown. Scale bar, 20 μm. *\*P* < 0.05, NS represents not significant (H_2_O_2_ groups versus 0 h group) (*n* = 50 cells). All data represent mean ± SD, and *P* values were established using Student’s *t*-test.

